# Frequency-resolved functional connectivity: Role of delay and the strength of connections

**DOI:** 10.1101/2020.09.10.291591

**Authors:** Abolfazl Ziaeemehr, Alireza Valizadeh

**Affiliations:** Department of Physics, Institute of Advanced Studies in Basic Sciences (IASBS), Zanjan, Iran

## Abstract

The brain functional network extracted from the BOLD signals reveals the correlated activity of the different brain regions, which is hypothesized to underlie the integration of the information across functionally specialized areas. Functional networks are not static and change over time and in different brain states, enabling the nervous system to engage and disengage different local areas in specific tasks on demand. Due to the low temporal resolution, however, BOLD signals do not allow the exploration of spectral properties of the brain dynamics over different frequency bands which are known to be important in cognitive processes. Recent studies using imaging tools with a high temporal resolution has made it possible to explore the correlation between the regions at multiple frequency bands. These studies introduce the frequency as a new dimension over which the functional networks change, enabling brain networks to transmit multiplex of information at any time. In this computational study, we explore the functional connectivity at different frequency ranges and highlight the role of the distance between the nodes in their correlation. We run the generalized Kuramoto model with delayed interactions on top of the brain’s connectome and show that how the transmission delay and the strength of the connections, affect the correlation between the pair of nodes over different frequency bands.

## 1 Introduction

A very prominent feature of brain networks is the ability to dynamically changing the routes for communication between the brain regions when undertaking different cognitive and executive functions (Park et al., 2018; Valdes-Sosa et al., 2011; Friston, 2011; Honey et al., 2007). This is revealed by extensive studies on the pattern of inter-relation between the activities of different brain regions at different brain states based on BOLD signals (Park et al., 2018; Allen et al., 2014; Calhoun et al., 2014; Chang and Glover, 2010; Wang et al., 2016). These correlated activities are supposed to underlie the integration of information over subsets of the whole-brain network, each comprising several regions (Friston, 2002). It is shown that each region can engage in one functional module and disengage from the other one due to the environmental demands and the state of the brain, enabling the brain to switch between multiplex of tasks across time. The recent advancement in the brain imaging using sophisticated EEG and MEG tools and developed methods of data analysis has made it possible to overcome the shortcomings of such approaches due to the high noise and difficulties in source localization (Haufe et al., 2011; Rodríguez-Rivera et al., 2006). The higher temporal resolution of these tools has extended the studies on the functional networks to the frequency domains which were not accessible through fMRI due to its low temporal resolution. This frequency range spans several specific bands which are believed to underlie different perceptional, cognitive, and executive functions, including delta, alpha, beta, and gamma bands (De Pasquale et al., 2010; Brookes et al., 2016; Li et al., 2017; Tewarie et al., 2016; Schnitzler and Gross, 2005). For example coherence in the gamma range is believed to provide a means for controlling effective communication between the brain regions (Bonnefond et al., 2017; Ray and Maunsell, 2015; Schroeder and Lakatos, 2009; Womelsdorf and Fries, 2007). Recent studies using MEG have shown that functional networks change in different frequency bands and multiplex of functional networks are present at any given time. These observations assert that any region can simultaneously participate in multiple functional modules, acting in parallel, and exploiting the structural communication channels for multiple tasks (Brookes et al., 2016). In this study, we question what properties of the brain structural network determine the pattern of the frequency-resolved functional network (Ziaeemehr et al., 2020a). Our focus is on the role of the distance between the nodes on the correction of their activity at different frequencies. Since both the delay in the interaction and the strength of the connections are dependent on the distance, we explore the effect of these two parameters by simulation of a simple model composed of phase oscillators, on top of the brain connectome network. Our results show that the dependence of the correlation on the distance is in general bolder at higher frequencies. We show that the variation of the correlation with frequency is faster for smaller delays, i.e., when the nodes are of shorter distance; while the connection strengths determine the amplitude of the variation of the correlation with frequency. Our results highlight the role of distance in the pattern of correlation between the nodes in brain networks.

## 2 Model

Our model consists of N phase oscillators each representing a region of interest of the brain and connected with time-delayed interactions described by a generalized Kuramoto model (Yeung and Strogatz, 1999):

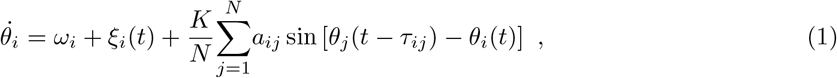

where, *θ*_*i*_ and *ω*_*i*_ = 2*πν*_*i*_ (*ν*_*i*_ being the frequency) are the phase and natural angular frequency of the *i*-th oscillator, respectively. *a*_*ij*_ are the elements of the adjacency matrix: *A. a*_*ij*_ = 1 if there is a link between the nodes *i* and *j* with a time delay *τ*_*ij*_; otherwise *a*_*ij*_ = 0. The parameter *K* sets the overall coupling strength. In our simulations, the initial values of *θ*_*i*_ are randomly drawn from a uniform distribution in the interval [0, 2*π*], and natural frequencies are drawn from a narrow normal distribution with given mean and variance. The degree of synchrony of the phase oscillators is quantified by the Kuramoto order parameters *r*, which is defined as 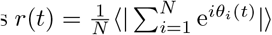. Here, ⟨…⟩ represents averaging over different network realizations and initial conditions. The magnitude is 0 ≤ *r* ≤ 1. The extreme cases are *r* = 1 (coherent state) and *r* = 0 (incoherent state). The time average of *r* after achieving a steady state is symbolized by *R*.

To measure the degree of synchronization between any two nodes of the network, we use the correlation index defined as *σ*_*ij*_ = ⟨cos[*θ*_*i*_(*t*) − *θ*_*j*_(*t*)] ⟩. Here *σ*_*ij*_ is an element of the correlation matrix *C* (Arenas et al., 2006).

The system of delayed differential equations (DDE) (Eq. 1) is solved numerically using adaptive Bogacki-Shampine (Flunkert, 2011) with minimum time step 0.05 [ms], absolute and relative error tolerance of 10^−8^ and 10^−5^, respectively. Noise can also be included in the differential equations, which are solved by the Euler-Maruyama method. For the simulations, we dropped the first 7 [s] and continued the simulations for 12 [s] and repeated the simulations 200 times with different initial conditions and frequency distribution. We also used a small amplitude of noise (0.05) and narrow normal distribution with a standard deviation of 0.01-0.1 for the initial frequencies. Adding small amplitude noise does not change the behavior of the system and only increases the transition time to a steady-state.

To quantify the pairwise distance between the distribution of values in functional networks and the connection strength of the connectome network, we used *pdist* module with the Euclidean metric from Scipy package (Oliphant, 2007).

We also considered a narrow interval of 0.05 to filter edges at a determined weight and an interval of 16 mm for selecting nodes with specific distance from each other. All the dropped units for time and distance are [ms] and [mm], respectively.

## 3 Results

In this paper, we aim to study the properties of the functional network of the brain at different frequency bands through simulation of a simple model of the human brain network. Specifically, we explore how the correlation between the nodes at different frequencies changes with the distance between the nodes. Our model is based on a generalized Kuramoto model run on top of the brain connectome composed of 66 nodes, whose properties are shown in Figure 1. The weight of the connections in the structural network, based on the number of axonal tracts between any two nodes is shown in Figure 1A, and the distance of the nodes is shown in Figure 1B. The structural network shows a modular structure at two levels, with 6 modules at the first level and two modules at the second (corresponding to two hemispheres). In Figure 1C and D we have shown the scatter plot of the connection strengths, and their mean and SD versus the distance between the nodes, respectively. In particular, it is seen that most strong connections are distributed around short distances with 2 < *d* < 5 cm, and distant nodes are only connected by weak links. Note that following other studies we assumed that the coupling strengths are scaled by the number of axonal fibers between any two nodes (Hagmann et al., 2008). When the connection strengths are then normalized, notably, 80% of them are very small < 10^−1^.

**Figure 1:**
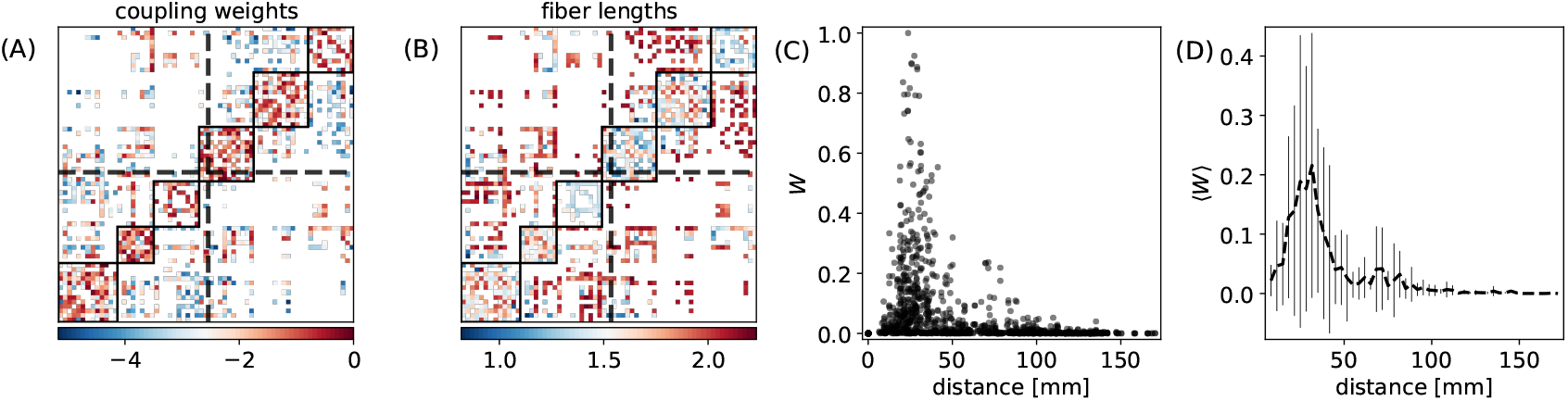
structural properties of the human connectome. (A) The normalized coupling weights and (B) length of fibers in the human connectome with 66 nodes (Hagmann et al., 2008). The solid squares show the modules and dash lines indicate the hemispheres. (C) Scatter plots of normalized coupling weights versus distance. (D) The distribution of mean the values of couplings and their standard deviation for different distances.

### 3.1 Frequency-resolved correlation matrix

In the model, we assume that the interaction between the nodes takes place through a delay time which in general is dependent on the distance. The distribution of the delays turns out to be the determinant factor for the functional network at different frequency bands. We first assume that the interaction delay is (linearly) proportional to the distance between the nodes, i.e., we take a fixed value for the speed of the signal transmission between the nodes (5 [m/s]). We also assume a weighted structural network where the connection strengths are scaled by the number of axonal tracts. The correlation matrix at five different frequency bands is shown in Fig. 2a. The appearance of anti-correlation between some pairs of nodes over higher frequency bands is apparent. Anti-correlation first appears between the nodes in different hemispheres over the beta range and in the gamma range, they are also observed for the intra-hemisphere pairs. This indicates the possible role of distance in the correlation between the nodes at different frequencies. The mean correlation between the nodes versus connection strength and distance is shown in Fig. 2b only for the pairs with a direct connection. At lower frequencies mean correlation shows no apparent conclusive dependence on the distance and connection strength. Anti-correlation appears at high distances at the beta range and shifts to lower distances with increasing frequency in accordance with the results shown in Fig. 2a. To get insight to the role of transmission delays and connection strength, we have shown scatter plots of the correlation between all the pairs at different frequencies in Fig 2c, where colors indicate the weights (left) and distances (right) between the nodes, respectively. It is seen that the mean correlation between all the pairs of nodes decreases at higher frequencies and negative correlation is observed at higher frequencies for distant nodes. The left panel shows that strong synapses lead to high positive and negative correlations at low and high frequencies, respectively. As it is seen in the right panel, at low frequencies the high positive correlation is seen mostly for low distances, while at higher frequencies short-distance nodes may show either positive or negative correlation with a high value. Long-distance nodes show lower values of correlation for all the frequencies. In the following, we inspect the frequency-resolved correlation matrix in more detail.

**Figure 2:**
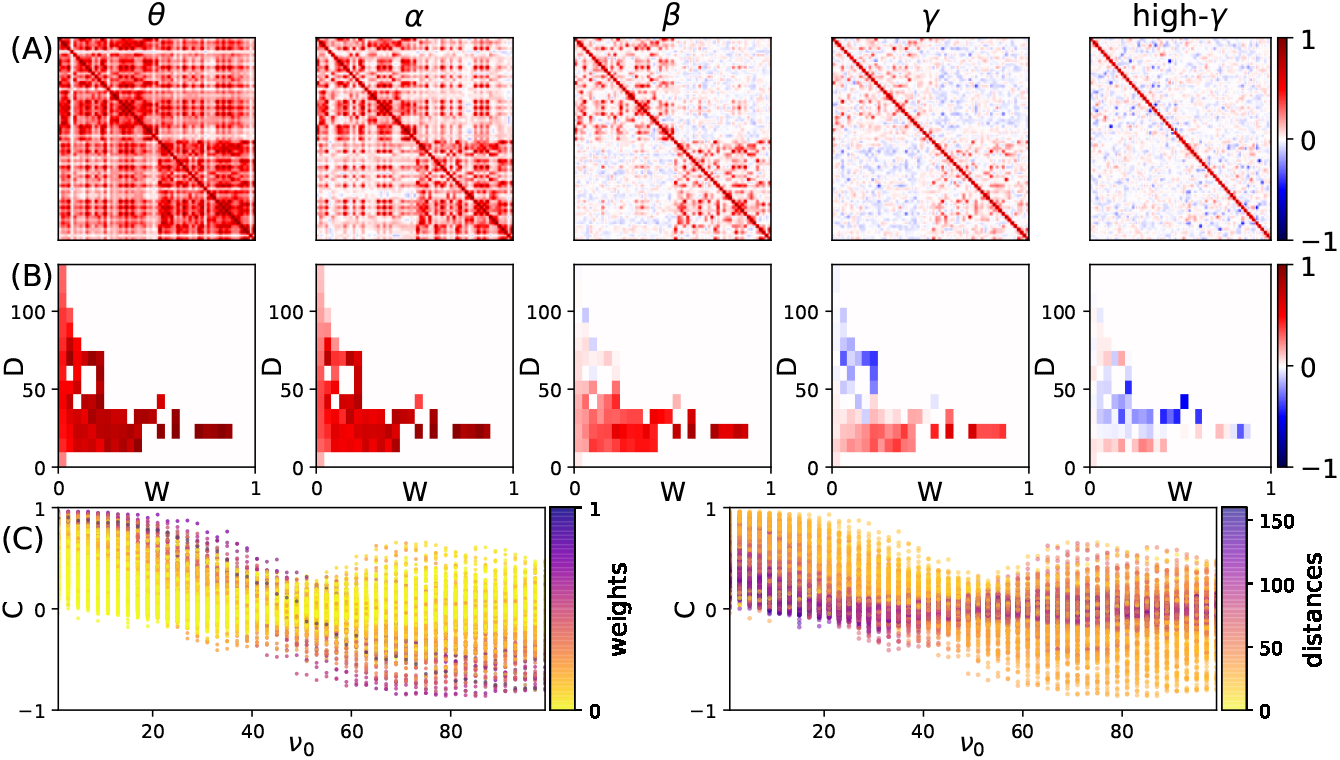
The correlation distributions. **(A)** The correlation matrices at five different frequencies. **(B)** The distribution of correlations versus weight and distance of connections at each frequency. **C** The distribution of correlations versus frequencies. The colors in left and right panels show the corresponding wights and distances, respectively.

### 3.2 Relation to distance and frequency

We have shown the scatter plot of the correlation of the pairs versus the distance of the nodes, at different frequencies in Fig. 3a. We observe a small negative correlation between the distance and correlation, i.e., those nodes which are farther from each other have a slightly lower correlation. But, while at lower frequencies the correlation almost linearly decreases with distance, at higher frequencies a steeper drop with distance is observed similar to the structural distribution of the connection weights. As an important corollary, we have compared this distribution with the structural one (Fig. 1c) by the distant measure introduced in Methods. The best similarity between the distribution of structural and dynamical couplings between the nodes (lowest distance between two distributions) is seen in beta and low gamma range around 30 Hz (Fig. 2b). We have also colored the points in Fig. 3a based on the weight of the structural link between the nodes. It is observed that strong links overall lead to a larger correlation between the nodes, but this is only observable at low distances since there are hardly strong links between the far nodes (Fig. 1c). Again, it is seen that while strong links lead to high positive correlation at low frequencies, they give rise to negative correlation at high-frequency ranges. To inspect more precisely, the relationship between the link strength and the correlation between the nodes, we have shown them in the scatter plots of Fig. 3c. Since in the connectome most of the links are very weak, the points in the scatter plot are packed in the small strength links. Nevertheless, the positive correlation between the link weight and correlation is observed for low frequencies and this correlation decreases in higher frequencies Fig. 3d. Notably, very weak links can carry high correlations in low frequencies and short distances (shown by color).

**Figure 3:**
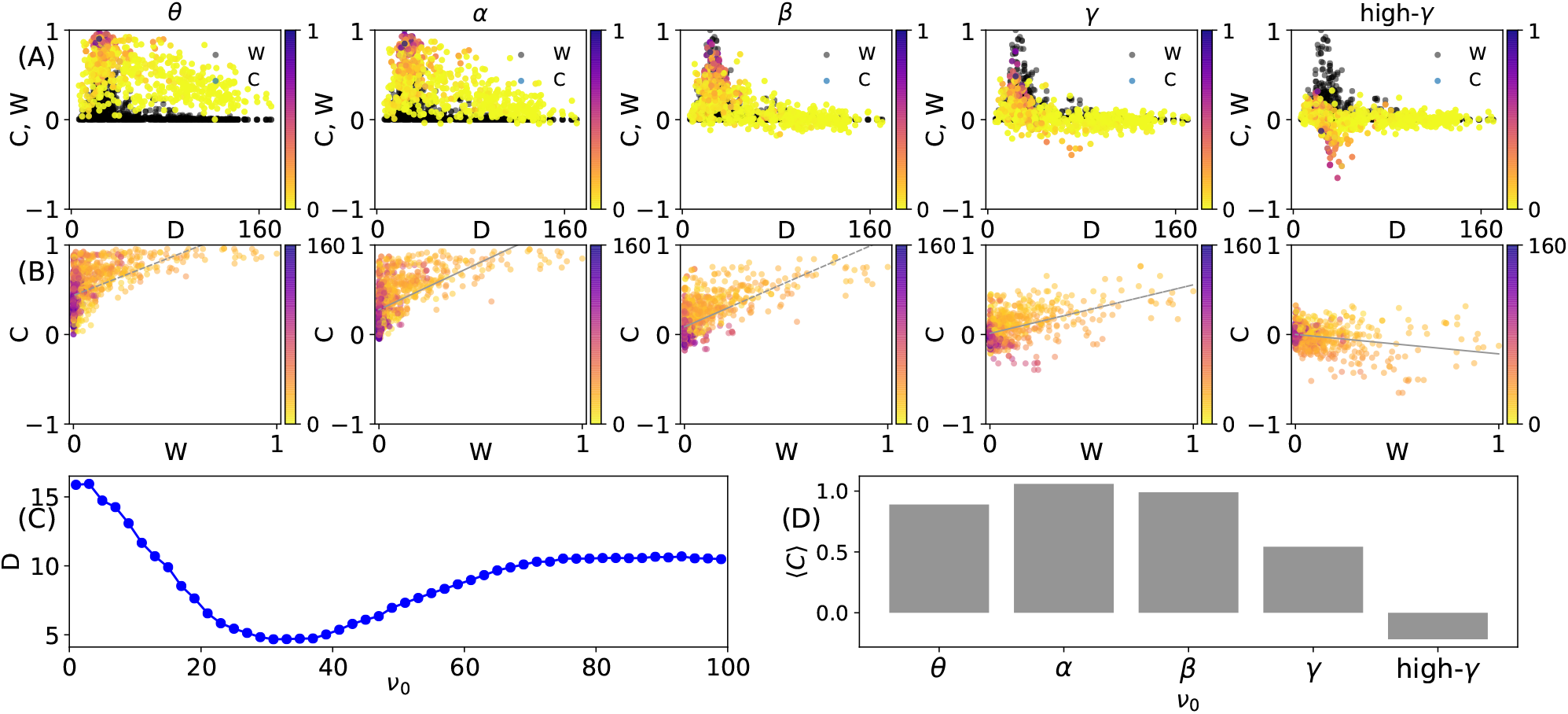
**(A)** The scatter plots of the correlations matrices versus distances. The colors indicate the corresponding weights of the connections. **(B)** The scatter plots of correlations matrices versus weights of connections. The colors indicate the corresponding distances at different frequencies. **(C)** The euclidean distance between scatter plots of correlations and weights of connections in the panels (A). **(D)** The slope of fitted lines in panel (B) versus frequency.

### 3.3 Distinct role of connection weight and delay

Since both the connection strength and the delay in communication between the nodes are dependent on the distance between the nodes, we question what is their distinct role on the pairwise correlation? More specifically, the results show that the distant nodes show smaller correlation at all frequency bands and they show anti-correlation for higher frequencies. Is that because they communicate through a longer delay or because they are connected by relatively weaker connections? To this end, we pick the pair of nodes with almost the same connection strength locating at different distances. Note that fixing the connection strength, only the delay is changing when the distance is varied. We have shown the mean correlation versus frequency for three different distances (delays) in Fig. 4a-b (for two different strengths). It can be seen that the correlation shows an almost periodic behavior with frequency and the variation in correlation is faster for the pairs with a longer delay. A comparison of the two panels also shows that the amplitude of the changes is larger for stronger connections, while the rate of the change with frequency is only dependent on the communication delay and is independent of the connection strengths.

**Figure 4:**
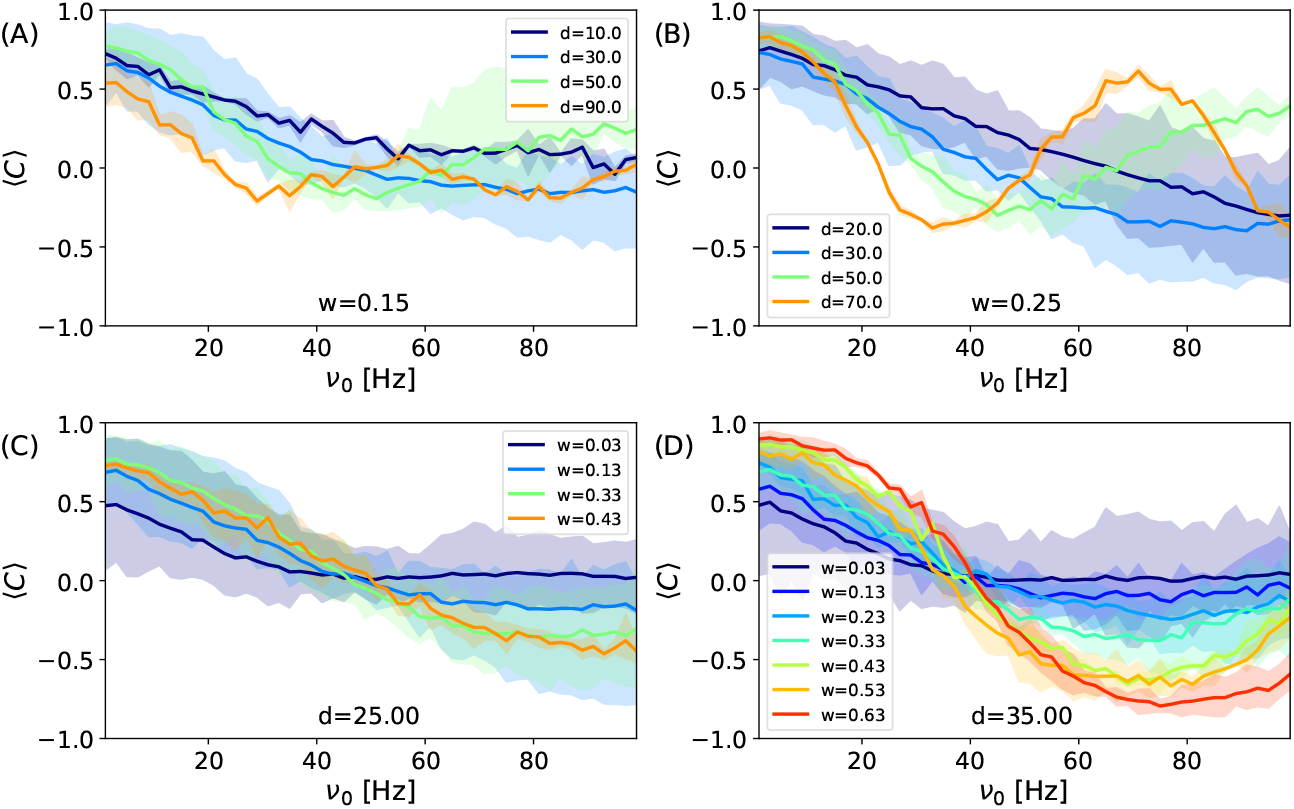
The correlation versus frequency for connections with fixed strength. **(A)** *W* ≈ 0.15 and **(B)** *W* ≈ 0.25 and various distances indicated in the legends. The correlation versus frequency for connections with fixed distance **(C)** *d* ≈ 25 mm and **(D)** *d* ≈ 35 mm and various strengths indicated in the legends. The colored areas show the results for p-value = 0.05.

To confirm the above results, we also presented the results for the nodes which are in almost the same distance but are connected by different connection strengths. The results presented in Fig. 4c-d (for two different distances) confirm that stronger connections lead to a larger amplitude of variation while for the pairs at the same distance, show the same rate of the change of correlation, with respect to frequency. Another point is that strong synapses not only give rise to higher correlation in low frequencies but also lead to more negative correlation at higher frequencies. To summarize, the presented results show that the rate of the changes in the correlation with frequency is determined by the transmission delay but the amplitude of the correlations is dependent on the link’s strength.

## 4 Discussion

In this manuscript, we studied the dependence of the correlation between the oscillatory activities of the pair of nodes to their distance, at different frequency bands, through simulation of a system of delayed-coupled phase oscillators on top of the brain’s connectome network. Since both delays in the communication between the nodes and the strength of the synaptic connections between them are a function of distance, we studied how the communication delay and connection strength can affect the correlation. We showed that the effect of these two parameters can be different at different frequencies. In particular, we found that at low frequencies the dependence of the correlation between the nodes is compatible with expectation and shorter delay and stronger connections lead to larger correlation. On the other hand at higher frequencies, the dependence is not trivial. Stronger connections in this range can lead to anti-correlation of the nodes and longer delays can both result in positive and negative correlation. In an intermediate-range, around beta and low gamma, we observed that the pattern of the correlations and the distribution of the weights against distance has maximal similarity to each other, compatible with the recent results (Ziaeemehr et al., 2020b).

Brain functional networks are constructed upon the statistical interdependencies between the activities of the brain regions which is conventionally measured by fMRI. The indirect measurement of the collective neuronal activity by fMRI can only reveal the slow dynamics of the brain due to its low temporal resolution, around one second (Sejnowski et al., 2014; Kim et al., 1997). Brain oscillations over several frequency bands which are known to be important for a variety of cognitive and executive functions have much shorter periods and it is impossible to assess them with BOLD signals. On the other hand, EEG and MEG recordings have a finer time resolution (Sejnowski et al., 2014; Burle et al., 2015) and recent instrumental advancements and improvements in data analysis software had made it possible to study frequency-resolved functional networks over wider frequency ranges(Hillebrand et al., 2012; Gramfort et al., 2013). These warrant the need for theoretical and computational studies on the spectral properties of the correlation matrix and the functional networks.

In the studies on the synchronization of the oscillators on complex networks, the connection strength and the interaction delays are two determinant factors which their effect is extensively explored (Cabral et al., 2012; Deco et al., 2009; Wang et al., 2014; Madadi Asl et al., 2018; Asl et al., 2018). It is shown the phase relations between the pair of the coupled oscillators depend on the connection strength and to the delay (Yeung and Strogatz, 1999; Sadeghi and Valizadeh, 2014; Esfahani et al., 2016; Esfahani and Valizadeh, 2014). Since these phase relations are hypothesized to underlie the communication between the brain populations, it is important to know how they change in realistic brain networks. In the brain networks, both the delay and connection strengths have a wide distribution making the brain structural network a very heterogeneous one. In this study, we used a realistic distribution for both the parameters and inspected how each of them impacts the pattern of the correlation between the brain regions, at different frequencies. With such a wide distribution of these parameters, a diversity of the correlations and the phase relations are observed which are important for a diverse and dynamic communication pattern in the brain (Maris et al., 2016; Ghosh et al., 2008; Breakspear et al., 2010).

While we did not directly explore the phase difference between the activities of the nodes, changes in the correlation could indirectly determine the phase relations. Namely, a high positive and negative correlation could indicate an almost in-phase or antiphase evolution of phases, respectively, with a continuum of intermediate phase differences between the two extremes. Our results indicated that the phase relations for any pair of nodes are in general dependent on the frequency. This has an important functional implication for the communication between the brain’s areas. Since the phase differences could determine the effective functional connectivity between the nodes (Maris et al., 2016; Friston, 2011), the pairs can communicate at different frequencies with different efficacy at multiple frequency bands. Such a multiplex of effective functional networks makes it possible to simultaneously engage the nodes at multiple functional modules (Park and Friston, 2013).

Moreover, our results showed more diverse phase relations at higher frequencies. Indeed over low-frequency bands, the correlation more slowly changes with distance and this means that long-range communication between the brain areas can take place by slow dynamics. On the other hand, a faster change in correlation with distance at high frequencies makes it possible to functionally dissociate the areas at a certain distance and form local functional modules. This can be a fundamental need for the brain networks for segregation of information processing at high-frequency bands and global integration at low frequencies (Isomura et al., 2006; Buzsáki and Mizuseki, 2014). The presence of multiple frequency bands could then lead to a hierarchy of spatial scales over which the information is integrated, corresponding to each frequency band (Zhou et al., 2006; Meunier et al., 2010). Our results show that the heterogeneous communication delay is the key requisite for the brain to enable such a hierarchical integration of information.

